## Supplemental Figures for "The organization of the gravity-sensing system in zebrafish"

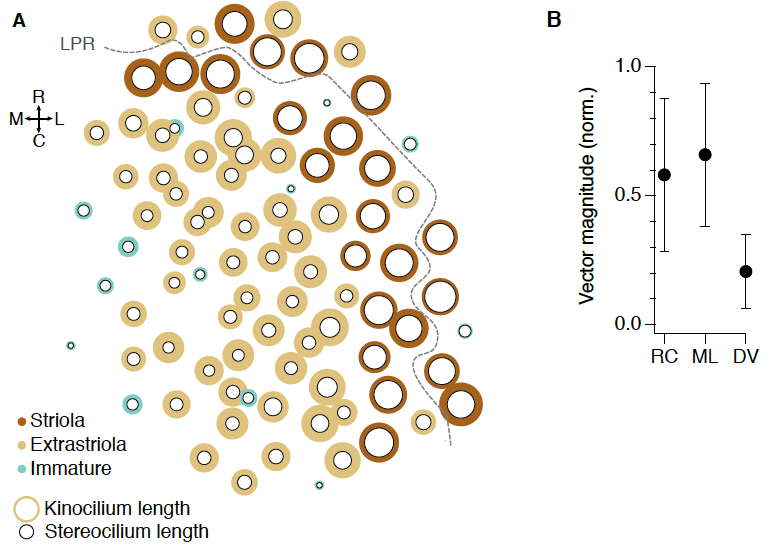

**Supplemental Fig. 1.** Hair cell ciliary lengths and tuning in different axes

**A,** Kinocilium and longest stereocilium lengths relative to the map of the utricular macula. Each hair cell is represented by an inner black circle indicating stereocilium length and an outer ring indicating kinocilium length, colorized as in Fig. 1. Hair cells whose stereocilium and kinocilium are similar in length are identified as striolar (brown), whereas those with a larger K/S ratio are identified as extrastriolar. Immature hair cells (blue-green) are identified by the very short lengths of their cilia.

**B,** The absolute vector magnitude extracted from hair cells in the rostrocaudal, mediolateral, and dorsoventral axes. Vectors were plotted in three dimensional space and normalized to a magnitude of 1; the magnitudes here are projections onto each axis (means ± SD). Dorsoventral tuning contributes little compared to the other two axes (Wilcoxon test,

ML vs DV: *p* = 9.7 x 10-21; DV vs RC: *p* = 9.7e-16; ML vs RC: *p* = 0.065).

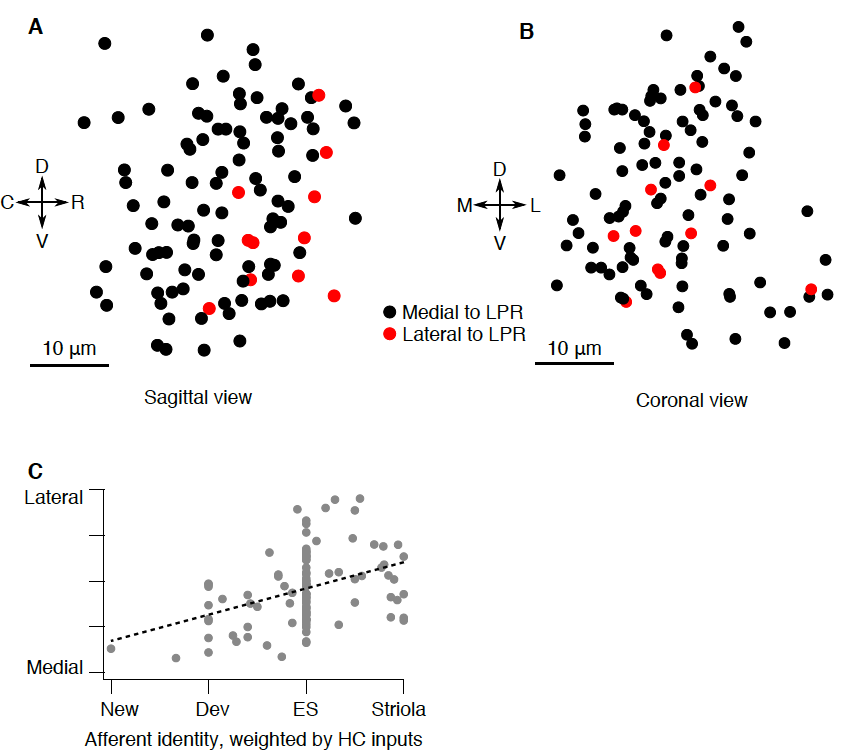

**Supplemental Fig. 2.** Utricular ganglion anatomy

**A,** Sagittal projection of the utricular ganglion showing the location of afferent somata that innervate hair cells medial (black) or lateral (red) to the line of polarity reversal. The afferents carrying contralateral head tilt information (i.e., innervating hair cells lateral to the LPR) are intermingled with the other afferents.

**B,** As in A, but coronal view.

**C,** Relationship between afferent soma position in the mediolateral axis and the hair cell types that innervate that afferent. Each afferent is assigned a value weighted by the number of ribbons it receives from hair cells with the given type. Significance of line fit, *p* = 3 x 10^-5^.

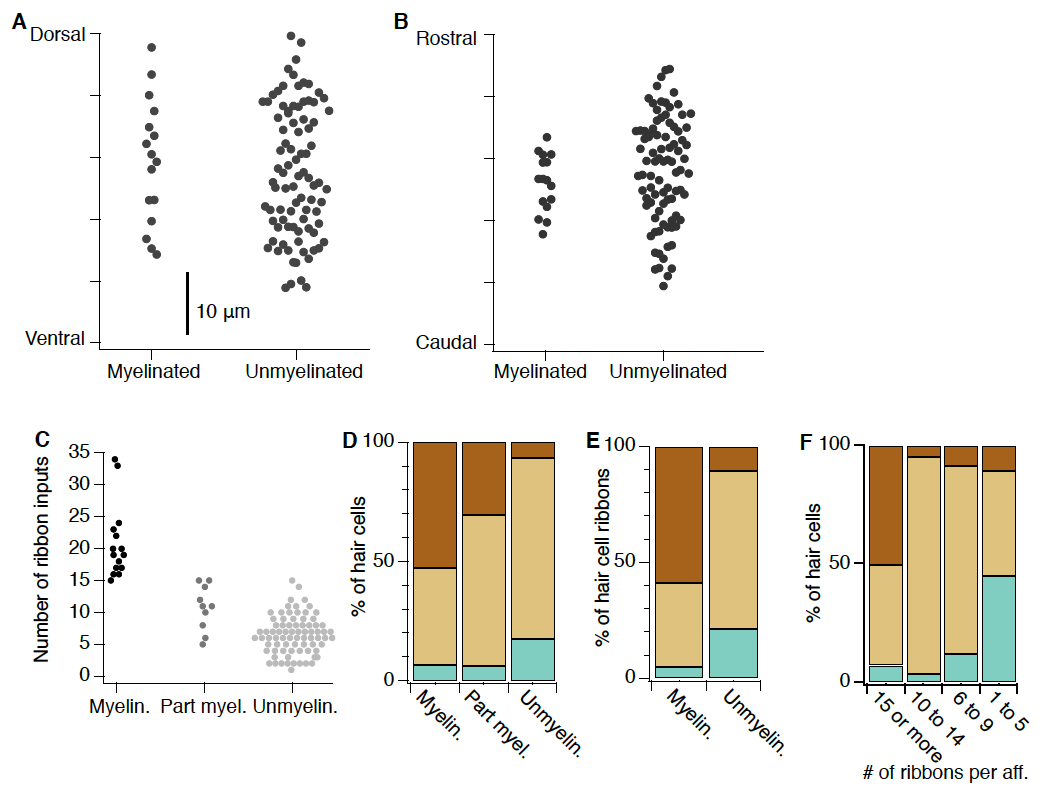

**Supplemental Fig. 3.** Ganglion developmental properties.

**A,** The dorsoventral soma position of utricular afferents that are or are not myelinated. There is no difference in position in this axis (*cf.* Fig. 4b).

**B,** As in A but for the rostrocaudal axis.

**C,** Number of ribbon synapses contacting each utricular afferent, as separated into those that appear fully myelinated, partially myelinated (myelination present on portions of the processes but absent for extended stretches), or wholly unmyelinated. Partially myelinated afferents exhibit properties intermediate to those of the myelinated and unmyelinated groups, with more ribbon inputs than wholly unmyelinated afferents (*p* = 0.0006, Wilcoxon) but fewer than fully myelinated afferents (*p* = 1.2 x 10^-6^, Wilcoxon).

**D,** Percent of hair cells with striolar (brown), extrastriolar (tan), or immature (blue-green) characteristics that are contacted by each afferent, separated into myelinated, partially myelinated, or unmyelinated as above. Again, partially myelinated afferents exhibit an intermediate phenotype to the other two groups.

**E,** Percent of hair cell ribbons contacted by myelinated or unmyelinated afferents, showing similar results as percent of hair cells (Fig. 4F).

**F,** Afferents with higher total ribbon contacts (≥ 15) preferentially receive inputs from striolar hair cells, whereas afferents with very few ribbon contacts (1-5) receive nearly half their input from immature hair cells.

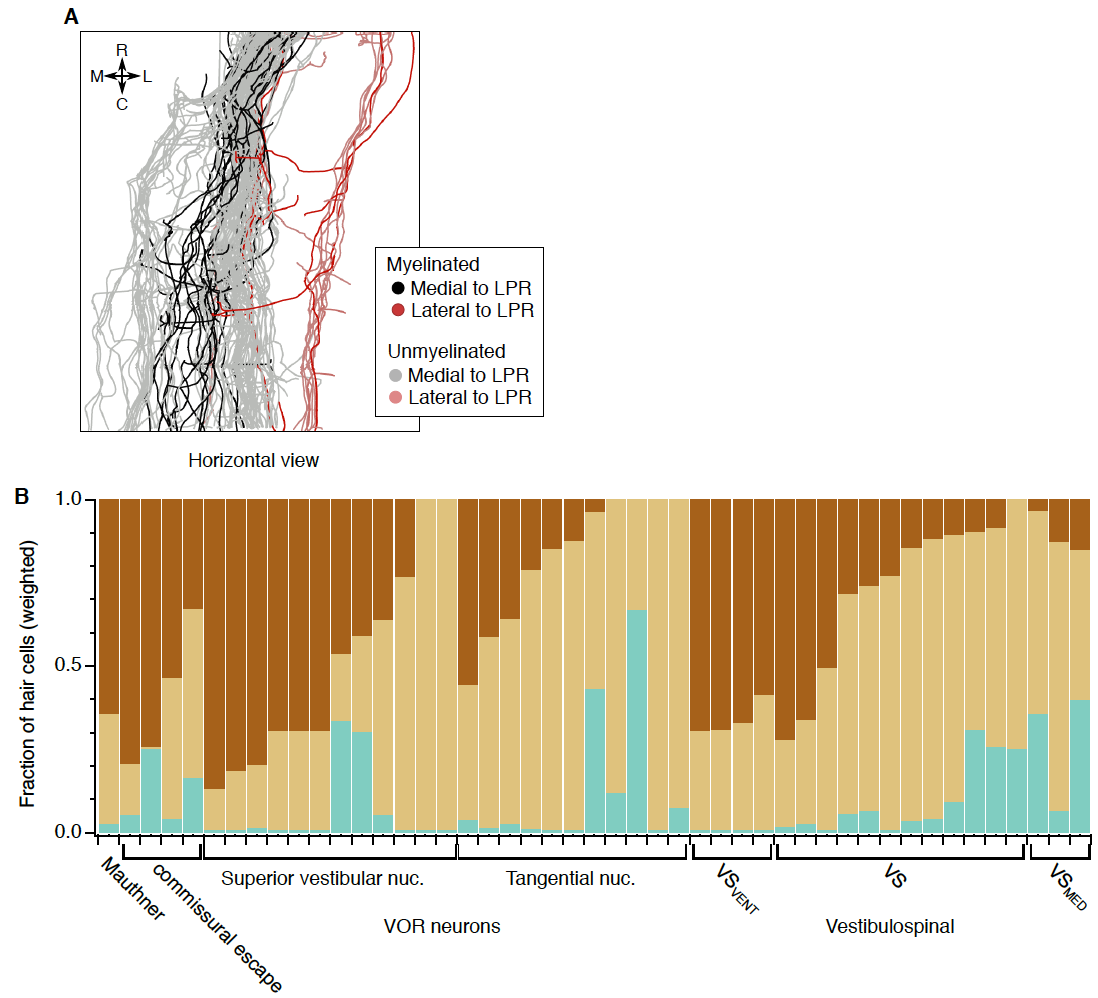

**Supplemental Fig. 4.** Central projections and connectivity.

**A,** Horizontal view of utricular afferent axons in the hindbrain. Axons are colorized by whether they are myelinated or unmyelinated (saturated vs tints) and whether they innervate hair cells medial or lateral to the line of polarity reversal (black vs red). The lateral-to-LPR afferents (red and pink), which encode head tilt to the contralateral side (= translation to the ipsilateral side), project much more laterally in the brainstem, and are largely separate from the majority of the afferents. In addition, there is some structure within the medial-innervating afferents, where myelinated axons run more centrally, in between two tranches of unmyelinated axons, suggesting a possible developmental growth pattern.

**B,** Weighted fraction of input to central neurons from striolar, extrastriolar, and immature hair cells, shown for every central neuron individually.

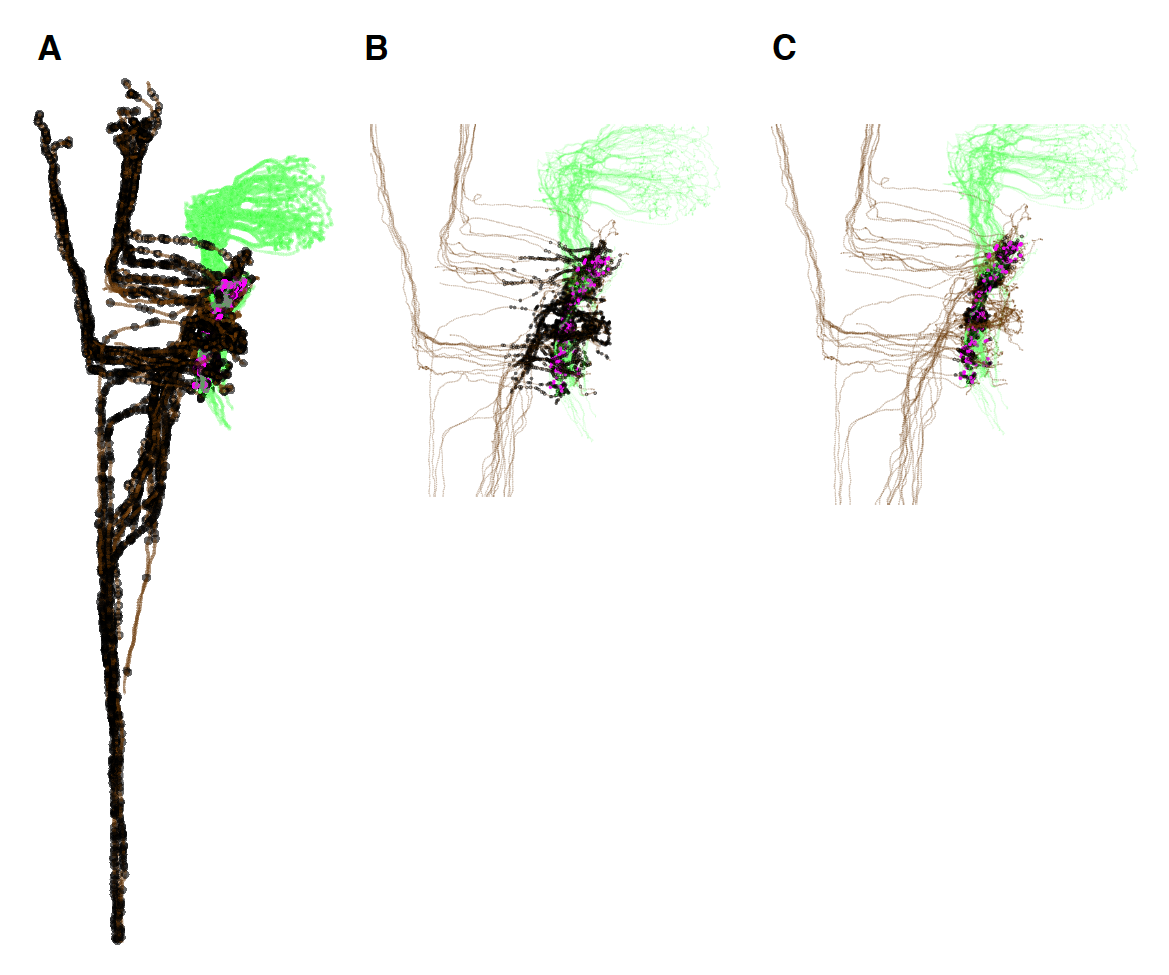

**Supplemental Fig. 5.** Representations of the Monte Carlo modeling approach.

**A,** Horizontal view of utricular circuits with the “unweighted” approach. Black circles indicate permissive areas for connectivity, whereas brown is neuronal processes without input.

**B,** Horizontal view of utricular circuits with the “50 µm within afferent” weighting approach.

**C,** Horizontal view of utricular circuits with the “5 µm within afferent” weighting approach.
